# Actin- and microtubule-based motors contribute to clathrin-independent endocytosis in yeast

**DOI:** 10.1101/2023.04.29.538819

**Authors:** Thaddeus K. Woodard, Daniel J. Rioux, Derek C. Prosser

## Abstract

Most eukaryotic cells utilize clathrin-mediated endocytosis as well as multiple clathrin-independent pathways to internalize proteins and membranes. Although clathrin-mediated endocytosis has been studied extensively and many machinery proteins have been identified, clathrin-independent pathways remain poorly characterized by comparison. We previously identified the first known yeast clathrin-independent endocytic pathway, which relies on the actin-modulating GTPase Rho1, the formin Bni1 and unbranched actin filaments, but does not require the clathrin coat or core clathrin machinery proteins. In this study, we sought to better understand clathrin-independent endocytosis in yeast by exploring the role of myosins as actin-based motors, since actin is required for endocytosis in yeast. We find that Myo2, which transports secretory vesicles, organelles and microtubules along actin cables to sites of polarized growth, participates in clathrin-independent endocytosis. Unexpectedly, the ability of Myo2 to transport microtubule plus ends to the cell cortex appears to be required for its role in clathrin-independent endocytosis. In addition, dynein, dynactin and proteins involved in cortical microtubule capture are also required. Thus, our results suggest that interplay between actin and microtubules contributes to clathrin-independent internalization in yeast.

**Summary Statement:** Clathrin-independent endocytosis is a poorly-understood but conserved process. Here, we provide evidence of a role for myosin and dynein as motor proteins involved the yeast clathrin-independent pathway.

## Introduction

In eukaryotic cells, the plasma membrane (PM) plays important roles in regulation of signal transduction, communication with the extracellular environment, and nutrient uptake. Maintenance of PM composition and function relies on a balance between exocytosis for delivery of membrane and proteins and on endocytosis for their internalization and removal from the cell surface. Exocytic and endocytic transport allow the cell to control the complement of proteins at the PM in order to amplify or attenuate responses to external cues, and permit selective uptake of nutrients by regulating surface availability of transporters. Additionally, endocytosis plays important quality control functions in mediating removal of damaged proteins through internalization and targeting to the lysosome for degradation.

Endocytic mechanisms have been studied extensively for over sixty years, where electron microscopy studies observed “bristle-coated” structures that were later identified as clathrin-coated pits and vesicles (Kanaseki and Kadota, 1969; Pearse, 1975; Roth and Porter, 1964). Subsequent studies have characterized clathrin-mediated endocytosis (CME) as the predominant pathway for internalization in most eukaryotic cells (Kaksonen and Roux, 2018). CME is highly conserved from yeast to human, and involves the action of more than 40 distinct proteins that arrive in a highly ordered sequence of events in order to select and concentrate cargo at sites of vesicle formation (Kaksonen et al., 2003; Kaksonen et al., 2005; Taylor et al., 2011). These proteins act in modules that correspond to discrete stages of clathrin-coated vesicle (CCV) formation, beginning with an initiation phase in which adaptor proteins bind to endocytic cargos at the PM as well as the assembling clathrin coat, effectively concentrating the cargo at a site of CCV formation (Howard et al., 2002; Maldonado-Báez et al., 2008; Newpher et al., 2005; Reider and Wendland, 2011; Reider et al., 2009). As the clathrin-coated site matures, endocytic accessory and scaffolding proteins facilitate additional clathrin assembly, and recruit proteins involved in later stages of vesicle formation, including actin-nucleating proteins in yeast required for the force generation needed for membrane invagination (Kaksonen et al., 2003; Kaksonen et al., 2005; Sun et al., 2006). In the late stages of CME, scission effectors such as dynamin and amphiphysins assist with membrane constriction at the neck of the budding vesicle, leading to separation of the CCV from the cell surface (Bliek et al., 1993; Damke et al., 1994; Kaksonen et al., 2005). Finally, uncoating of the CCV permits recycling of endocytic machinery proteins and fusion of the vesicle with downstream compartments (Ungewickell et al., 1995). Studies have shown that the modular nature of CME is well conserved, and genetic or pharmacological perturbation of any stage in the process can reduce the efficiency of cargo internalization (Goode et al., 2015; Kaksonen and Roux, 2018).

Early studies of endocytosis also observed vesicle formation in the absence of a bristle coat (Anderson and Batten, 1983; Morris and Saelinger, 1983), suggesting that multiple mechanisms for internalization likely existed. Indeed, numerous clathrin-independent endocytic (CIE) pathways have been identified across eukaryotes. These include a variety of mechanistically distinct routes of internalization. For example, CIE encompasses phagocytic and macropinocytic pathways that rely on membrane protrusion or ruffling, lipid (typically cholesterol)-enriched membrane microdomains such as caveolae and clathrin-independent carriers and GPI-enriched endocytic compartments (CLIC/GEEC), and actin-dependent pathways relying on Arf-and Rho-family small GTPases (Howes et al., 2010; Lamaze et al., 2001; Mayor et al., 2014; Radhakrishna et al., 1996; Sabharanjak et al., 2002; Sharma et al., 2002). In addition, recent studies have demonstrated that ultrafast endocytosis at synaptic terminals is clathrin-independent, and fast endophilin-mediated endocytosis (FEME) is a distinct CIE pathway that relies on Endophilin A1, dynein and microtubules for internalization of PM proteins (Boucrot et al., 2015; Casamento and Boucrot, 2020; Watanabe et al., 2013a; Watanabe et al., 2013b). Despite the variety of CIE pathways that have been documented, our current understanding of the molecular mechanisms governing any CIE pathway is poor in comparison to that of CME. Reasons for this disparity may include the comparative lack of cargos that utilize CIE pathways for entry, and a lack of genetically tractable model systems for studies of CIE.

We previously identified the first-known *Saccharomyces cerevisiae* CIE pathway using a mutant strain lacking four monomeric clathrin-binding adaptor proteins: the epsins Ent1 and Ent2, and the AP180/PICALM homologs Yap1801 and Yap1802 (Prosser et al., 2011). Yeast lacking these adaptors (*ent1*Δ *ent2*Δ *yap1801*Δ *yap1802*Δ, also known as 4Δ) have defective CME as seen by retention of endocytic cargos at the PM, while any one full-length adaptor is sufficient for endocytosis (Maldonado-Báez et al., 2008). Although *ENT1* and *ENT2* constitute an essential gene pair, expression of the phosphatidylinositol (4,5)-bisphosphate [PI(4,5)P_2_]- binding epsin N-terminal homology (ENTH domain) of either gene is sufficient for viability but not for their role in CME (Aguilar et al., 2006; Maldonado-Báez et al., 2008; Wendland et al., 1999). Thus, 4Δ+ENTH1 cells expressing the ENTH domain of Ent1 have been a useful model for studying deficits in CME (Maldonado-Báez et al., 2008; Prosser et al., 2011). Using this strain to identify genes whose overexpression restored endocytic cargo internalization, we found that high-copy expression of the cell wall stress sensor *MID2*, the Rho1 guanine nucleotide exchange factor *ROM1* and the actin-modulating small GTPase *RHO1* all enhanced internalization of multiple endocytic cargos, and that their function did not require clathrin or the major CME machinery proteins (Prosser and Wendland, 2012; Prosser et al., 2011). The budding yeast CIE pathway additionally requires the formin Bni1, which generates unbranched actin filaments independent of Arp2/3, proteins in the polarisome complex which recruits and activates Bni1 at sites of polarized growth, and proteins that stabilize unbranched actin filaments such as the tropomyosins Tpm1 and Tpm2. Further studies identified α-arrestins as cargo-selective proteins that participate in CIE through mechanisms that are distinct from their established roles in CME, as well as a dual role for the early-acting CME protein Syp1, suggesting that some proteins may contribute to multiple endocytic pathways (Apel et al., 2017; Prosser et al., 2015).

Although yeast were thought to rely solely upon CME prior to our identification of the Rho1-dependent CIE pathway, other studies have also suggested that yeast do not strictly require CME for cargo internalization. For example, *Candida albicans* can perform endocytosis in the absence of functional clathrin and Arp2/3 (Epp et al., 2010; Epp et al., 2013), while endocytosis at the cylindrical sides of cells in the fission yeast *Schizosaccharomyces pombe* appears to depend on the formin For3 (Gachet and Hyams, 2005). Overall, our mechanistic understanding of CIE pathways in yeast, and indeed in any eukaryotic cell type, lags considerably behind that of CME due at least in part to a lack of tools to characterize these pathways.

In this study, we extend our earlier findings by expanding the set of proteins required for CIE in yeast. Since all endocytosis in yeast critically depends on actin polymerization, we specifically focused on roles for myosins as actin-based motors and on myosin-interacting proteins. We find that Myo2, which transports organelles and other cellular structures along actin cables to sites of polarized growth, is required for CIE. Unexpectedly, Myo2-dependent transport of microtubule plus ends, as well as dynein, dynactin, and proteins involved in cortical microtubule capture participate in CIE, while transport of other structures is dispensable. Thus, interplay between actin and microtubule cytoskeletons may play important roles in yeast CIE.

## Results

### The type V myosin Myo2 is required for clathrin-independent endocytosis

Our previous studies of CIE demonstrated a requirement for the formin Bni1, which localizes mainly to the bud tip and bud neck, where it promotes actin elongation at the barbed end of unbranched filaments (Evangelista et al., 2001; Prosser et al., 2011; Pruyne et al., 2002; Sagot et al., 2002). Additionally, we found that the tropomyosin Tpm1, which stabilizes unbranched actin filaments, was necessary for CIE. These filaments are bundled into actin cables that serve as tracks for myosin-dependent delivery of material to sites of polarized growth (Pruyne et al., 1998). Our initial findings underscored the role of actin in CIE, and prompted us to examine the contribution of myosins as actin-based motors. Budding yeast possess five myosins: the type I myosins Myo3 and Myo5 are involved in CME, the type II myosin Myo1 is involved in constriction of the cytokinetic ring at the bud neck, and the type V myosins Myo2 and Myo4 transport proteins, vesicles, organelles, and mRNA along actin cables (Bobola et al., 1996; Geli and Riezman, 1996; Pruyne et al., 2004; Watts et al., 1987). Of these, Myo2 appears to play the most prominent role in transport along unbranched actin cables; thus, we focused on this protein as a candidate motor involved in CIE.

*MYO2* is an essential gene, and thus cannot be deleted (Johnston et al., 1991). Instead, we generated *myo2* mutants with reduced processivity (Schott et al., 1999; Schott et al., 2002). The lever arms of Myo2 permit movement along actin filaments, and contain IQ repeats that can be truncated to generate motors that take shorter “steps”. Using full-length (6IQ) or truncated (4IQ or 2IQ) mutants expressed as the sole source of Myo2 in WT, 4Δ+Ent1 (with functional CME) and 4Δ+ENTH1 (with impaired CME) backgrounds, we examined localization of the pheromone receptor Ste3-GFP as an endocytic cargo. Ste3 is a seven-transmembrane receptor that is normally transported to the PM in haploid *MATα* cells, constitutively internalized via CME, and delivered to the vacuole for degradation (Davis et al., 1993). Under conditions where CME is blocked, Ste3 is instead largely retained at the PM, with reduced transport to the vacuole (Maldonado-Báez et al., 2008). We previously found that expression of *ROM1* from a high-copy plasmid promoted clathrin-independent internalization of Ste3-GFP in numerous CME-deficient mutants including 4Δ+ENTH1 (Prosser et al., 2011); thus, we asked whether a reduction in Myo2 processivity would impair *ROM1*-dependent activation of CIE. In WT and 4Δ+Ent1 backgrounds transformed with empty vector, expression of Myo2^6IQ^, Myo2^4IQ^, or Myo2^2IQ^ resulted in Ste3-GFP localization primarily at the vacuole (Fig. 1A). Thus, reduced Myo2 processivity does not appear to affect CME. In contrast, Ste3-GFP showed increased retention at the PM in 4Δ+ENTH1 cells transformed with empty vector and expressing full-length and truncated Myo2, which is consistent with defective endocytosis in 4Δ+ENTH1 cells. When these 4Δ+ENTH1 cells were instead transformed with high-copy *ROM1* to promote CIE, we found that Ste3-GFP internalization and transport to the vacuole was improved in cells expressing full-length Myo2^6IQ^, but showed less or no improvement in cells expressing Myo2^4IQ^ or Myo2^2IQ^ compared to equivalent empty vector-transformed cells.

**Figure 1:**
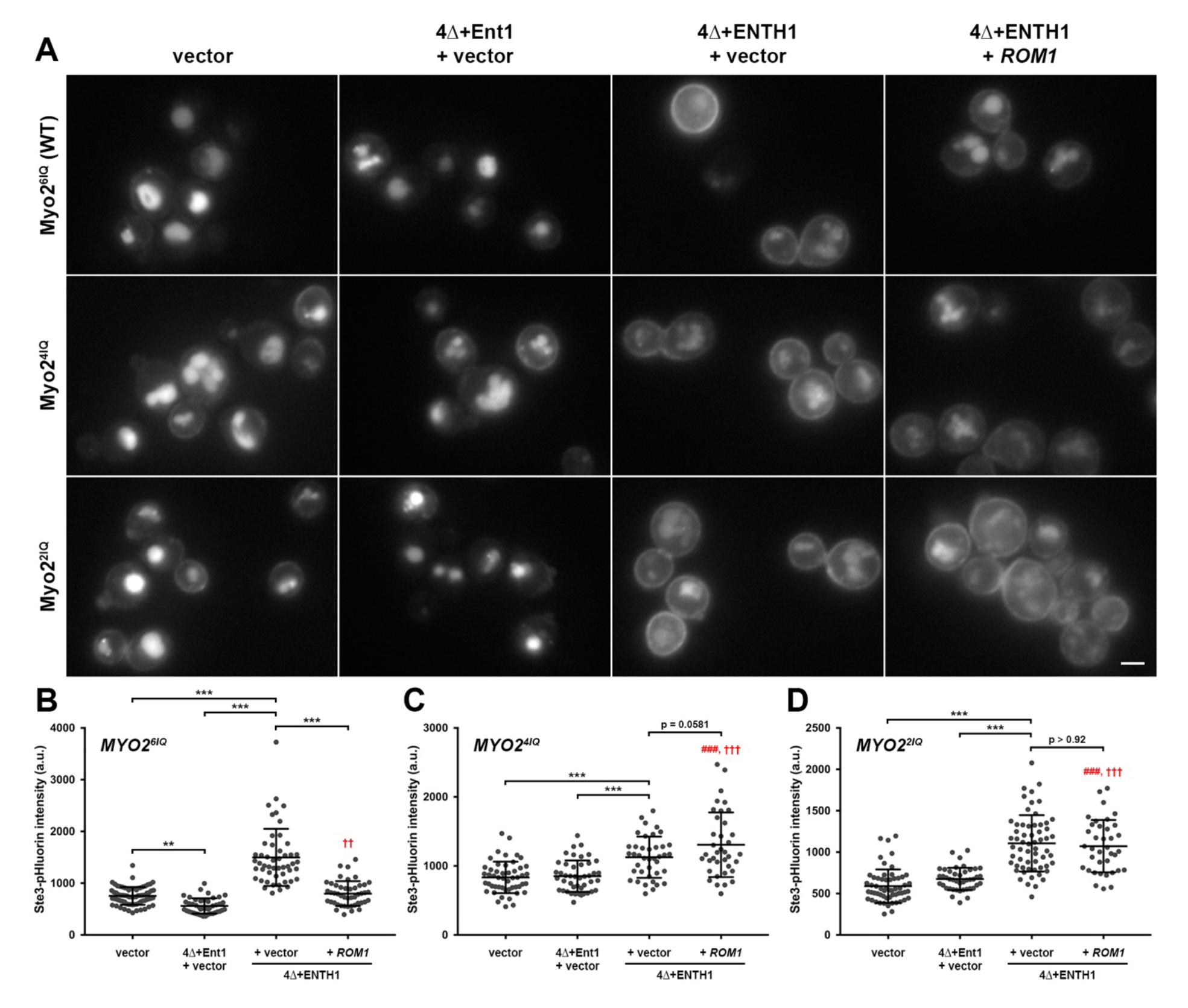
Effect of Myo2 IQ repeat truncation on clathrin-independent endocytosis. Full-length Myo2^6IQ^ or truncated Myo2^4IQ^ or Myo2^2IQ^ mutants expressed as the sole source of Myo2 in WT (vector), 4Δ+Ent1 and 4Δ+ENTH1 backgrounds were transformed with vector or high-copy *ROM1* as indicated. (A) Cells expressing Ste3-GFP from the endogenous locus were imaged by fluorescence microscopy. (B-D) Quantification of whole cell fluorescence intensity in cells expressing Ste3-pHluorin transformed as in *A*. (B) Myo2^6IQ^ (n=73, n=45, n=50 and n=46, respectively). (C) Myo2^4IQ^ (n=50, n=44, n=40 and n=37, respectively). (D) Myo2^2IQ^ (n=58, n=43, n=60 and n=37, respectively). Mean ± s.d.; \*\*\**P*<0.001; ###*P*<0.001 compared to WT; ††*P*<0.01 and †††*P*<0.001 compared to 4Δ+Ent1). Scale bar: 2 µm.

We previously developed a quantitative method for assessing endocytic capacity in live yeast cells by tagging the cytoplasmic tail of cargo proteins with superecliptic pHluorin, a pH-sensitive variant of GFP (Miesenböck et al., 1998; Prosser et al., 2010; Prosser et al., 2016; Sankaranarayanan et al., 2000). When the tail of the cargo is exposed to a neutral environment such as the cytoplasm, the pHluorin tag is brightly fluorescent. In contrast, fluorescence is quenched upon packaging of the tag into acidic environments, such as the intraluminal vesicles within multivesicular bodies or the vacuole lumen. Measurement of steady-state, whole-cell Ste3-pHluorin intensity can thus reveal changes in endocytosis, as cells that efficiently internalize Ste3-pHluorin and target it to the vacuole are dim, while cells with defective endocytosis retain the tagged protein at the PM and are comparatively bright (Prosser et al., 2010; Prosser et al., 2016). Using the same truncated Myo2 strains expressing genomically-tagged Ste3-pHluorin instead of GFP, we were able to quantitatively assess the effect of reduced Myo2 processivity on CIE. As expected for cells with defective CME, Ste3-pHluorin intensity in empty vector-transformed 4Δ+ENTH1 cells expressing full-length Myo2^6IQ^ or truncated Myo2^4IQ^ or Myo2^2IQ^ was significantly brighter than in the equivalent WT and 4Δ+Ent1 backgrounds (Fig. 1B-D). In Myo2^6IQ^-expressing 4Δ+ENTH1 cells, transformation with high-copy *ROM1* partially restored Ste3-pHluorin internalization, as shown by a significant reduction in fluorescence intensity compared to the same cells with empty vector (Fig. 1B). In contrast, high-copy *ROM1* failed to improve Ste3-pHluorin internalization in 4Δ+ENTH1 cells expressing Myo2^4IQ^ (Fig. 1C) or Myo2^2IQ^ (Fig. 1D), as these cells remained as bright as, or brighter than, equivalent empty vector-transformed cells. Taken together, these data suggest that Myo2 processivity is required for CIE in yeast.

To examine the potential contribution of other myosins to CIE, we generated single *myo1*Δ and *myo4*Δ deletions in WT, 4Δ+Ent1 and 4Δ+ENTH1 backgrounds expressing Ste3-GFP. Both *myo1*Δ and *myo4*Δ did not alter vacuolar Ste3-GFP localization in WT and 4Δ+Ent1 backgrounds transformed with empty vector, suggesting that neither gene is required for cargo internalization when CME is functional (Supplementary Fig. S1A-B). Whereas Ste3-GFP was partially retained at the PM in empty vector-transformed *myo1*Δ or *myo4*Δ 4Δ+ENTH1 cells, transformation with high-copy *ROM1* reduced cell surface retention and increased vacuole localization in both cases, suggesting that neither Myo1 nor Myo4 is required for CIE.

Since Myo3 and Myo5 play overlapping roles in force generation at cortical actin patches in yeast, deletion of both genes simultaneously is required to block CME (Geli and Riezman, 1996). Consistent with this finding, *myo3*Δ *myo5*Δ cells transformed with empty vector showed a severe retention of Ste3-GFP at the plasma membrane, with cargo fluorescence virtually undetectable in the vacuole (Supplementary Fig. S1C-D). This defect could be complemented by low-copy expression of *MYO5*, which restored vacuolar Ste3-GFP localization and reduced cell surface retention of the cargo. In contrast, high-copy *YAP1801*, which restored endocytosis in 4Δ+ENTH1 cells, had no effect on cargo internalization in *myo3*Δ *myo5*Δ (Prosser et al., 2011). Although we predicted that high-copy *ROM1* would promote CIE in *myo3*Δ *myo5*Δ cells since these myosins are known to function in CME, we unexpectedly found that Ste3-GFP was strongly retained at the PM. However, close examination of these cells revealed dim vacuolar fluorescence with high-copy *ROM1*, but not in cells transformed with empty vector or *YAP1801* (Supplementary Fig. 1C-D). The limited ability of Rom1 to promote Ste3 internalization in *myo3*Δ *myo5*Δ cells might be explained by previous studies showing that actin cable morphology is severely altered in this strain (Anderson et al., 1998; Goodson et al., 1996). Indeed, phalloidin staining of *myo3*Δ *myo5*Δ cells with empty vector revealed numerous actin patches, but no obvious actin cable structures (Supplementary Fig. 1E). While low-copy *MYO5* restored actin cables, they remained absent in cells with high-copy *YAP1801* or *ROM1*. Thus, Myo3 and Myo5 may play roles in CIE, but their role might be explained by an indirect effect on actin cable morphology. Taken together, these data suggest that Myo2 processivity contributes to CIE, while other myosins may play less prominent roles.

### Myo2-dependent transport of microtubules is required for CIE

Previous studies showed that Myo2 plays a critical role in mother-to-bud transport of secretory vesicles, mitochondria, vacuoles, peroxisomes, and microtubules (Beach et al., 2000; Boldogh et al., 2004; Fagarasanu et al., 2005; Fagarasanu et al., 2009; Ishikawa et al., 2003; Pruyne et al., 1998; Schott et al., 1999). To achieve this, the cargo-binding domain (CBD) of Myo2 interacts with organelle-specific adaptors that link each structure to the motor for transport (Eves et al., 2012; Fagarasanu et al., 2009; Jin et al., 2011; Lipatova et al., 2008; Pashkova et al., 2006). Structure-function analysis of the CBD revealed unique surface patches that associate with Myo2 adaptors, and mutations within these patches result in selective loss of adaptor binding and organelle transport (Eves et al., 2012; Pashkova et al., 2006). To assess which functions are necessary for CIE, we expressed CBD mutants from the endogenous *MYO2* locus as the sole source of Myo2 in WT, 4Δ+Ent1 and 4Δ+ENTH1 backgrounds. We focused on five point mutants, each defective in one transport function: D1297N (loss of Vac17 binding and vacuole transport); K1312A (loss of Mmr1 binding and mitochondrial transport); K1408A (loss of Kar9 binding and microtubule plus end transport); Q1447R (deficient in binding the Rab GTPases Ypt31, Ypt32, Ypt11 and Sec4, reduction in secretory vesicle transport); and E1484A (loss of Inp2 binding and peroxisome transport; (Eves et al., 2012; Fagarasanu et al., 2009; Ishikawa et al., 2003; Lipatova et al., 2008; Pashkova et al., 2006)). As seen in Fig. 2A, expression of any of these mutants in WT or 4Δ+Ent1 cells transformed with empty vector did not alter vacuolar localization of Ste3-GFP compared to cells with wild-type Myo2, suggesting that none of these Myo2 functions impact CME. As expected for 4Δ+ENTH1 cells expressing Myo2 CBD mutants and transformed with empty vector, Ste3-GFP showed retention at the PM. Expression of high-copy *ROM1* improved Ste3-GFP internalization and vacuolar delivery in 4Δ+ENTH1 cells expressing wild-type Myo2 and all of the CBD mutants except for except for the microtubule transport-deficient mutant, Myo2^K1408A^.

**Figure 2:**
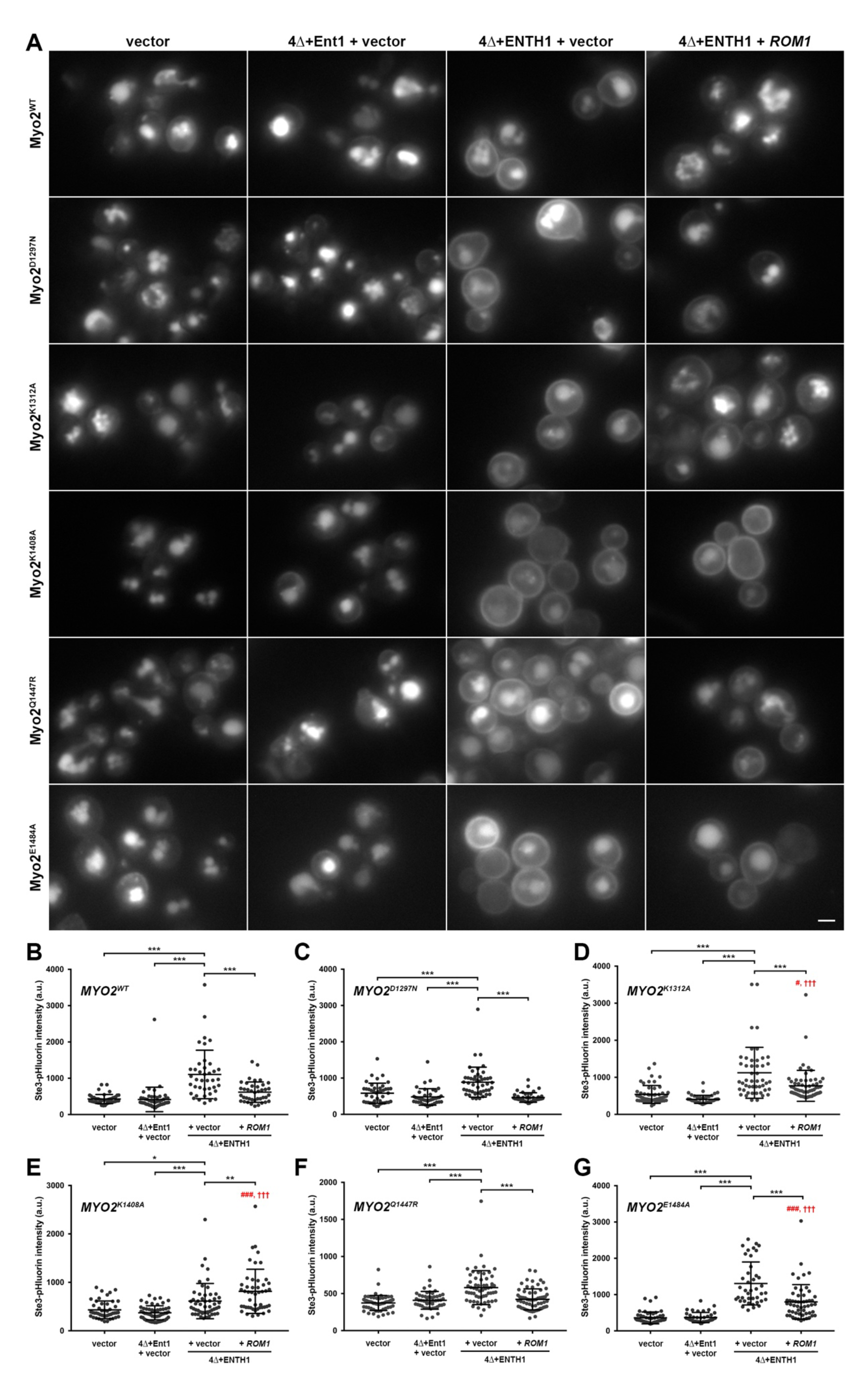
Effect of Myo2 cargo-binding domain mutation on clathrin-independent endocytosis. Wild-type Myo2 or CBD mutants (D1297N, K1312A, K1408A, Q1447R, and E1484A) expressed as the sole source of Myo2 in WT (vector), 4Δ+Ent1 and 4Δ+ENTH1 backgrounds were transformed with vector or high-copy *ROM1* as indicated. (A) Cells expressing Ste3-GFP from the endogenous locus were imaged by fluorescence microscopy. (B-G) Quantification of whole cell fluorescence intensity in cells expressing Ste3-pHluorin transformed as in *A*. (B) Myo2^WT^ (n=46, n=52, n=40 and n=40, respectively). (C) Myo2^D1297N^ (n=43, n=41, n=49 and n=41, respectively). (D) Myo2^K1312A^ (n=53, n=47, n=48 and n=63, respectively). (E) Myo2^K1408A^ (n=46, n=55, n=52 and n=48, respectively). (F) Myo2^Q1447R^ (n=59, n=52, n=58 and n=65, respectively). (G) Myo2^E1484A^ (n=51, n=41, n=41 and n=54, respectively). Mean ± s.d.; \**P*<0.05, \*\**P*<0.01, \*\*\**P*<0.001; #*P*<0.05 and ###*P*<0.001 compared to WT; †††*P*<0.001 compared to 4Δ+Ent1). Scale bar: 2 µm.

When we analyzed endocytic capacity in the same set of strains expressing Ste3-pHluorin, we observed quantitative trends that agreed with localization patterns of Ste3-GFP (Fig. 2B-G). In WT and 4Δ+Ent1 backgrounds with functional CME, whole-cell intensity of Ste3-pHluorin was similarly low regardless of whether the cells expressed wild-type Myo2 or any of the CBD mutants. Ste3-pHluorin intensity was significantly higher in all empty vector-transformed 4Δ+ENTH1 strains compared to their Myo2 CBD-matched control WT and 4Δ+Ent1 background strains, as expected for cells with defective CME. Transformation of 4Δ+ENTH1 cells expressing wild-type Myo2 or the D1297N, K1312A, Q1447R, or E1484A CBD mutants significantly improved Ste3-pHluorin internalization, as seen by lower fluorescence intensity compared to empty vector-transformed 4Δ+ENTH1 cells (Fig. 2B-D and F-G). In contrast, Myo2^K1408A^ 4Δ+ENTH1 cells transformed with high-copy *ROM1* were significantly *brighter* than empty vector-transformed cells, suggesting a possible worsening of the endocytic defect (Fig. 2E). Interestingly, high-copy *ROM1* showed varying degrees of rescue in the other CBD mutants: it fully rescued Ste3-pHluorin internalization to WT or 4Δ+Ent1 levels in cells expressing Myo2 WT, D1297N and Q1447R, but only partially improved internalization to levels that remained higher than for WT or 4Δ+Ent1 in cells expressing Myo2^K1312A^ or Myo2^E1484A^. Overall, the ability of Myo2 to transport microtubule plus ends appears to be required for CIE, while other motor functions appear to be dispensable.

### Links between Myo2 and microtubule plus ends are required for CIE

Our finding that microtubule plus end transport contributes to CIE was unexpected because the yeast microtubule cytoskeleton is thought to play limited roles in membrane traffic; however, a previous study implicated microtubules and the plus-end tracking protein Bik1 in trafficking of the v-SNARE Snc1 (Boscheron et al., 2016). *tub1-Glu* and *bik1*Δ mutations caused Snc1 mislocalization reminiscent of defective endocytosis that were suppressed by Rho1 activation; thus, we further examined links between Myo2 and microtubule plus end-binding proteins. The Myo2 CBD binds Kar9, the yeast homolog of adenomatous polyposis coli (APC; (Beach et al., 2000)). In turn, Kar9 associates with the plus end-binding protein Bim1, which is homologous to mammalian EB1. Myo2, Kar9 and Bim1 function together in a pathway required for cortical microtubule (cMT) capture which is partially redundant with a second pathway involving dynein, Bik1 (homolog of CLIP-170) and Num1 (Adames and Cooper, 2000; Farkasovsky and Küntzel, 2001; Lee et al., 2000; Miller and Rose, 1998; Miller et al., 2000; Yin et al., 2000). Kar9 and Bim1 both additionally associate with Bik1, suggesting the possibility of cross-talk between these pathways (Moore et al., 2006).

To begin examining the role of these plus end-binding proteins in CIE, we generated *kar9*Δ, *bim1*Δ and *bik1*Δ mutants in WT, 4Δ+Ent1 and 4Δ+ENTH1 backgrounds expressing Ste3-GFP or Ste3-pHluorin. Each of the plus end transport-regulating proteins did not alter vacuolar delivery of Ste3-GFP in empty vector-transformed WT and 4Δ+Ent1 backgrounds compared to equivalent cells with no modification to plus end-binding genes (Fig. 3A). As expected, Ste3-GFP was partially retained at the PM in 4Δ+ENTH1 cells as well as *kar9*Δ, *bim1*Δ, and *bik1*Δ 4Δ+ENTH1 cells with empty vector. High-copy *ROM1* appeared to improve Ste3-GFP internalization in 4Δ+ENTH1 and *bik1*Δ 4Δ+ENTH1 cells, but had little effect in *kar9*Δ or *bim1*Δ 4Δ+ENTH1 cells. When we quantified Ste3-pHluorin intensity in these backgrounds, we found that WT and 4Δ+Ent1 backgrounds were similarly dim in each case, while 4Δ+ENTH1 backgrounds were significantly brighter in comparison (Fig. 3B-E). In 4Δ+ENTH1 cells with no modification to plus-end binding genes, high-copy *ROM1* fully restored Ste3-pHluorin internalization to WT or 4Δ+Ent1 levels (Fig. 3B). In contrast, high-copy *ROM1* did not reduce Ste3-pHluorin intensity in *kar9*Δ 4Δ+ENTH1 cells compared to empty vector, suggesting that the plus end-binding complex is required for CIE (Fig. 3C). Ste3-pHluorin internalization was partially restored in *bim1*Δ and *bik1*Δ 4Δ+ENTH1 cells with high-copy *ROM1*, where fluorescence intensity was significantly lower than cells with empty vector, suggesting that neither gene is required for CIE (Fig. 3D-E). This was unexpected, as *ROM1* appeared to improve Ste3-GFP internalization in *bik1*Δ, but not *bim1*Δ 4Δ+ENTH1 cells (Fig. 3A). Notably, the decrease in Ste3-pHluorin intensity for *bim1*Δ 4Δ+ENTH1 cells expressing high-copy *ROM1* versus empty vector (14.6% reduction) was much smaller than that of *bik1*Δ 4Δ+ENTH1 cells (47.4% reduction), suggesting a greater effect for *bim1*Δ than for *bik1*Δ. Combined with the PM retention of Ste3-GFP in *bim1*Δ 4Δ+ENTH1 cells, it is thus possible that Bik1 plays a weaker role in CIE, or that the two proteins share overlapping function through their function in distinct cMT capture pathways or interactions with Kar9. Unfortunately, *bim1*Δ *bik1*Δ cells are inviable, so we were unable to directly test this possibility (Schwartz et al., 1997). Nonetheless, these data suggest that Kar9 and microtubule plus end-binding proteins contribute to CIE.

**Figure 3:**
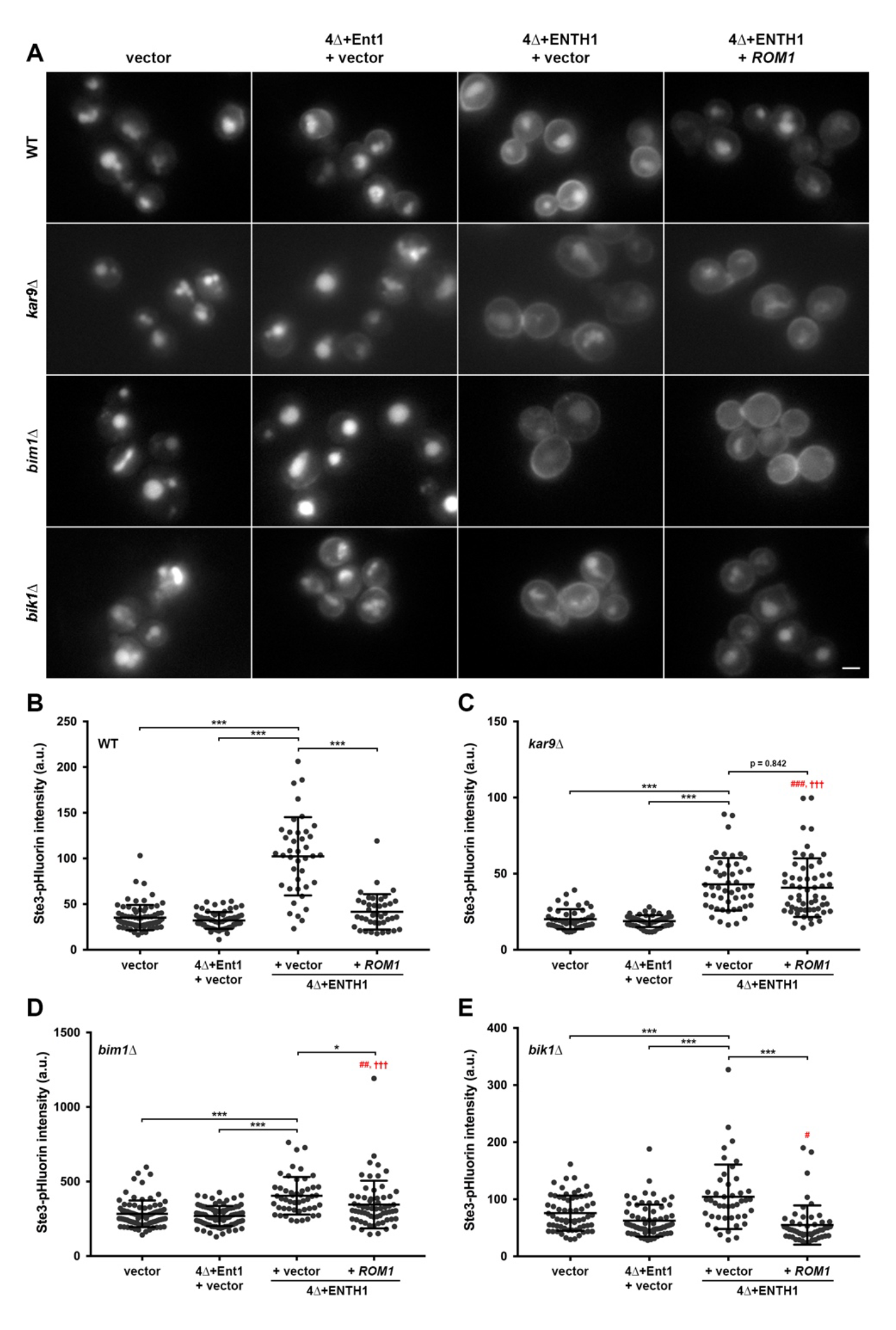
Requirement for microtubule plus end-binding proteins in clathrin-independent endocytosis. WT (vector), 4Δ+Ent1 and 4Δ+ENTH1 backgrounds were transformed with vector or high-copy *ROM1* as indicated. *kar9*Δ, *bim1*Δ and *bik1*Δ were generated in the same strains. (A) Cells expressing Ste3-GFP from the endogenous locus were imaged by fluorescence microscopy. (B-E) Quantification of whole cell fluorescence intensity in cells expressing Ste3-pHluorin transformed as in *A*. (B) WT, 4Δ+Ent1 and 4Δ+ENTH1 backgrounds with no additional modifications (n=77, n=65, n=40 and n=40, respectively). (C) *kar9*Δ (n=45, n=51, n=50 and n=57, respectively). (D) *bim1*Δ (n=84, n=84, n=50 and n=63, respectively). (E) *bik1*Δ (n=64, n=70, n=44 and n=52, respectively). Mean ± s.d.; \**P*<0.05, \*\*\**P*<0.001; #*P*<0.05, ##*P*<0.01 and ###*P*<0.001 compared to WT; †††*P*<0.001 compared to 4Δ+Ent1). Scale bar: 2 µm.

### Stability of cytoplasmic microtubules contributes to CIE

Given our observations that Myo2-and Kar9-dependent transport of microtubules is required for CIE in yeast, we decided to more directly test the involvement of microtubules in this process. We previously showed that actin is required for CIE in addition to its established role in CME by treating cells with the actin-depolymerizing drug Latrunculin A (LatA; (Prosser et al., 2011)). Using WT, 4Δ+Ent1 and 4Δ+ENTH1 cells transformed with empty vector, as well as 4Δ+ENTH1 cells with high-copy *ROM1*, we examined Ste3-GFP localization in vehicle (DMSO), LatA, and nocodazole-treated cells to determine whether microtubule depolymerization would also affect Rom1-dependent internalization (Fig. 4). As seen previously, two-hour treatment with LatA potently inhibited endocytosis in all strains, where Ste3-GFP was almost completely retained at the PM (Prosser et al., 2011). In contrast, DMSO-treated controls showed the expected pattern of primarily vacuolar Ste3-GFP in WT and 4Δ+Ent1 cells with empty vector, prominent PM retention in 4Δ+ENTH1 cells with empty vector, and improved internalization in 4Δ+ENTH1 with high-copy *ROM1*. Ste3-GFP showed similar localization in nocodazole-and DMSO-treatment conditions for WT, 4Δ+Ent1 and 4Δ+ENTH1 cells with empty vector. In contrast, 4Δ+ENTH1 cells with high-copy *ROM1* appeared to show weaker Ste3-GFP internalization with nocodazole-treatment compared to DMSO. Thus, microtubules appear to participate in CIE.

**Figure 4:**
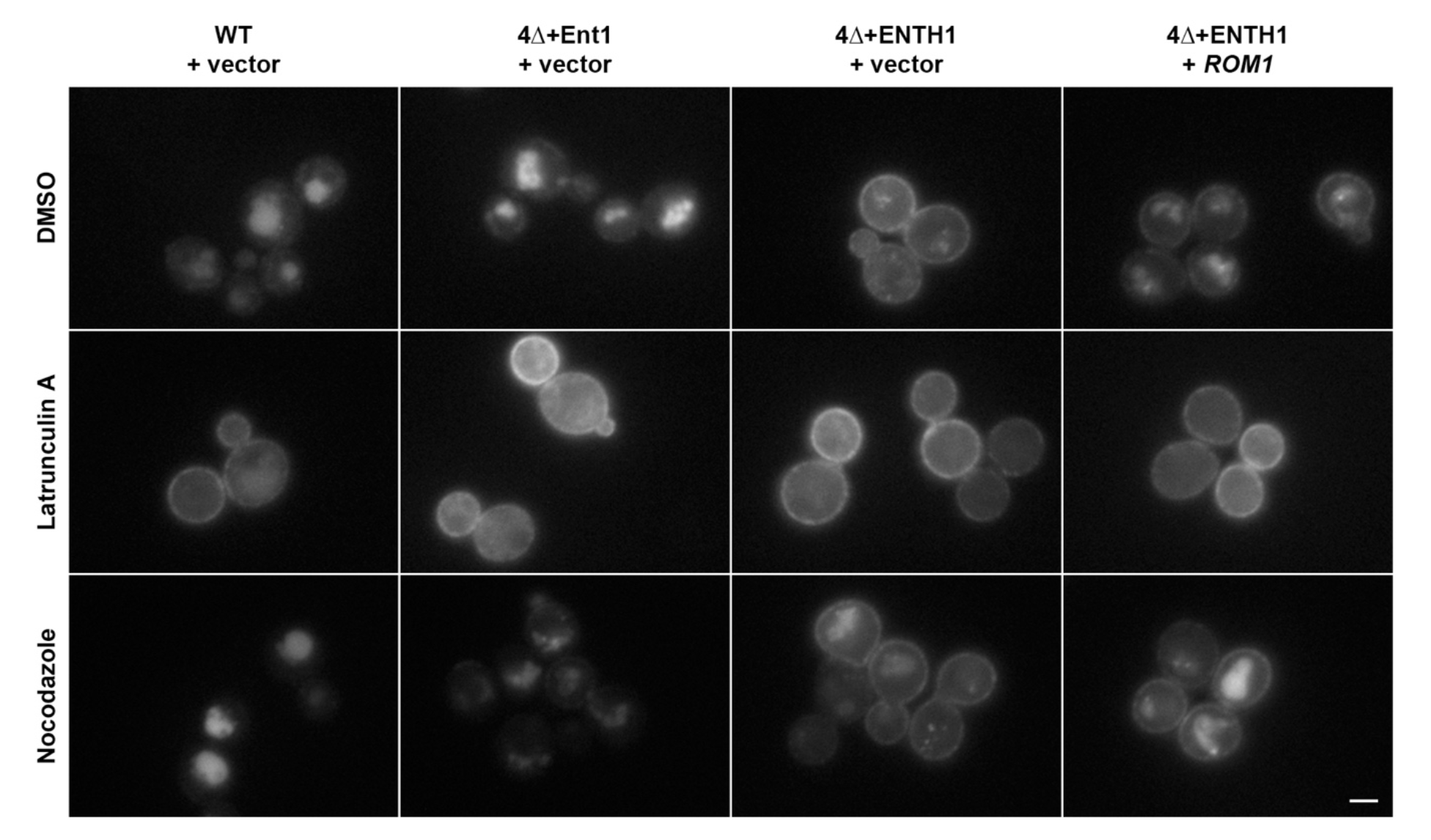
Requirement for actin and microtubules in clathrin-independent endocytosis. WT, 4Δ+Ent1 and 4Δ+ENTH1 cells expressing Ste3-GFP from the endogenous locus were transformed with vector or high-copy *ROM1* as indicated. Cells grown to mid-logarithmic phase were treated with vehicle (DMSO), 200 µM Latrunculin A, or 15 µg/ml Nocodazole for 2 h prior to imaging by fluorescence microscopy. Scale bar: 2 µm.

Since force must be exerted at the PM for endocytosis to occur, we further reasoned that stability of cytoplasmic microtubules is required for CIE. The kinesin-related motors Kip2 and Kip3 play opposing roles in regulating cytoplasmic microtubule stabilization, where Kip2 inhibits catastrophe and promotes microtubule polymerization, while Kip3 destabilizes microtubules (Miller et al., 1998). Thus, *kip2*Δ results in destabilization of cytoplasmic microtubules, while *kip3*Δ causes their hyper-stabilization. We generated *kip2*Δ and *kip3*Δ strains in WT, 4Δ+Ent1 and 4Δ+ENTH1 backgrounds, and used these to examine Ste3-GFP localization. Neither *kip2*Δ nor *kip3*Δ altered the predominantly vacuolar localization of Ste3-GFP in WT or 4Δ+Ent1 backgrounds with empty vector, as expected for strains with functional CME (Fig. 5A). In *kip2*Δ 4Δ+ENTH1 cells with empty vector, Ste3-GFP largely accumulated at the PM, consistent with a block in CME. Transformation of *kip2*Δ 4Δ+ENTH1 cells with high-copy *ROM1* did not noticeably improve Ste3-GFP internalization. Interestingly, empty vector-transformed *kip3*Δ 4Δ+ENTH1 cells showed lower PM accumulation of Ste3-GFP with correspondingly higher vacuolar delivery, and high-copy *ROM1* gave similar distribution of the cargo. These findings correlate with microtubule stability: we were unable to observe cytoplasmic microtubules in *kip2*Δ cells expressing GFP-Tub1 compared to WT cells; in contrast, *kip3*Δ cells showed exaggerated cytoplasmic microtubules that appeared to wrap around the cell cortex (Fig. 5B).

**Figure 5:**
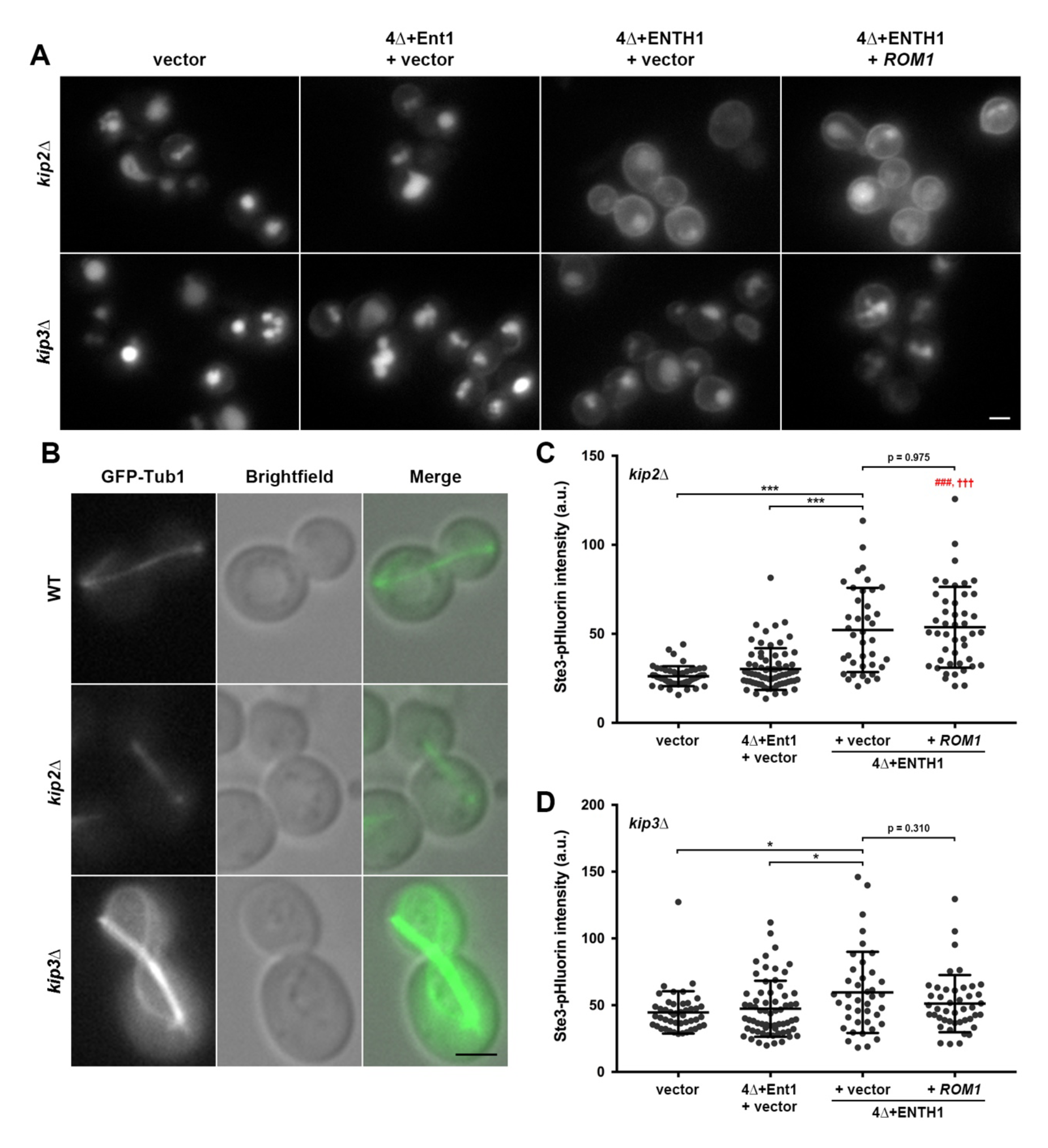
Role of cytoplasmic microtubule stability in clathrin-independent endocytosis. Cytoplasmic microtubule-destabilizing (*kip2*Δ) and -stabilizing (*kip3*Δ) mutants generated in WT (vector), 4Δ+Ent1 and 4Δ+ENTH1 backgrounds were transformed with vector or high-copy *ROM1* as indicated. (A) Cells expressing Ste3-GFP from the endogenous locus were imaged by fluorescence microscopy. (B) Fluorescence microscopy of GFP-Tub1 expressed in large-budded WT, *kip2*Δ and *kip3*Δ cells, with brightfield and merged panels to demonstrate microtubule localization. (C-D) Quantification of whole cell fluorescence intensity in cells expressing Ste3-pHluorin transformed as in *A*. (C) *kip2*Δ (n=49, n=69, n=39 and n=45, respectively). (D) *kip3*Δ (n=46, n=65, n=40 and n=44, respectively). Mean ± s.d.; \**P*<0.05, \*\*\**P*<0.001; ###*P*<0.001 compared to WT; †††*P*<0.001 compared to 4Δ+Ent1). Scale bars: 2 µm.

Quantification of Ste3-pHluorin intensity in *kip2*Δ and *kip3*Δ strains supported our observations of Ste3-GFP localization. For *kip2*Δ or *kip2*Δ 4Δ+Ent1 cells with empty vector, whole-cell Ste3-pHluorin intensity was similarly low, while *kip2*Δ 4Δ+ENTH1 cells with empty vector were significantly brighter (Fig. 5C). Consistent with a requirement for Kip2 in CIE, transformation of *kip2*Δ 4Δ+ENTH1 cells with high-copy *ROM1* failed to improve internalization, as shown by similar brightness to *kip2*Δ 4Δ+ENTH1 cells with empty vector. Deletion of *KIP3* in 4Δ+ENTH1 cells with empty vector also resulted in a small, but significant increase in Ste3-pHluorin intensity compared to *kip3*Δ or *kip3*Δ 4Δ+Ent1 cells (Fig. 5D); however, this increase (33.7% for *kip3*Δ 4Δ+ENTH1 compared to *kip3*Δ with empty vector) was much smaller than seen when comparing *kip2*Δ to *kip2*Δ 4Δ+ENTH1 with empty vector (99.2% increase). Transformation of *kip3*Δ 4Δ+ENTH1 cells with high-copy *ROM1* resulted in Ste3-pHluorin intensity levels that were indistinguishable from *kip3*Δ, *kip3*Δ 4Δ+Ent1, or *kip3*Δ 4Δ+ENTH1 cells with empty vector. Taken together, these experiments demonstrate that stability of cytoplasmic microtubules is required for CIE, and that their stabilization may increase clathrin-independent endocytosis in yeast.

### Dynein and dynactin are required for CIE in yeast

Our findings that cytoplasmic microtubules, and Myo2-dependent transport of microtubule plus ends, contribute to CIE prompted us next to examine whether the minus end-directed microtubule motor dynein and its cofactor dynactin are also needed. As microtubule plus ends are transported along actin cables, dynein and dynactin are targeted to the plus end through an interaction between the dynein heavy chain (Dyn1) and Pac1 (homolog of the mammalian lissencephaly protein, LIS1), which in turn binds to Bik1 (Sheeman et al., 2003). To explore the role of this protein module in CIE, we focused on four deletions: *dyn1*Δ (dynein heavy chain) and *pac11*Δ (dynein intermediate chain) which cause a loss of dynein function, *nip100*Δ (dynactin p150^Glued^ subunit) which causes a loss of dynactin function, and *pac1*Δ which disrupts plus end-tracking of dynein/dynactin (Geiser et al., 1997; Sheeman et al., 2003). We generated each of these deletions in WT, 4Δ+Ent1, and 4Δ+ENTH1 backgrounds, and subsequently examined Ste3-GFP localization and the effect of high-copy *ROM1* on Ste3 internalization in the CME-defective 4Δ+ENTH1 strain. As expected for each WT and 4Δ+Ent1 background transformed with empty vector, Ste3-GFP was efficiently delivered to the vacuole, indicating that loss of dynein/dynactin function or plus end-tracking are not required for cargo internalization when CME is functional (Fig. 6A). In *dyn1*Δ, *pac11*Δ, *nip100*Δ, and *pac1*Δ 4Δ+ENTH1 cells transformed with empty vector, Ste3-GFP localized prominently at the PM, indicating a reduction in internalization. When each of the corresponding 4Δ+ENTH1 backgrounds were instead transformed with high-copy *ROM1*, Ste3-GFP retention at the PM was similar to cells with empty vector. Quantification of Ste3-pHluorin intensity confirmed the endocytic effects observed with Ste3-GFP: *dyn1*Δ, *pac11*Δ, *nip100*Δ, and *pac1*Δ in WT and 4Δ+Ent1 backgrounds with empty vector had similarly low fluorescence intensity, which was significantly higher in the 4Δ+ENTH1 background (Fig. 6B-E). Transformation of *dyn1*Δ, *pac11*Δ, *nip100*Δ, and *pac1*Δ 4Δ+ENTH1 cells with high-copy *ROM1* failed to reduce the elevated Ste3-pHluorin intensity, indicating that Rom1 was no longer able to improve cargo internalization in the absence of dynein/dynactin or Pac1 function.

**Figure 6:**
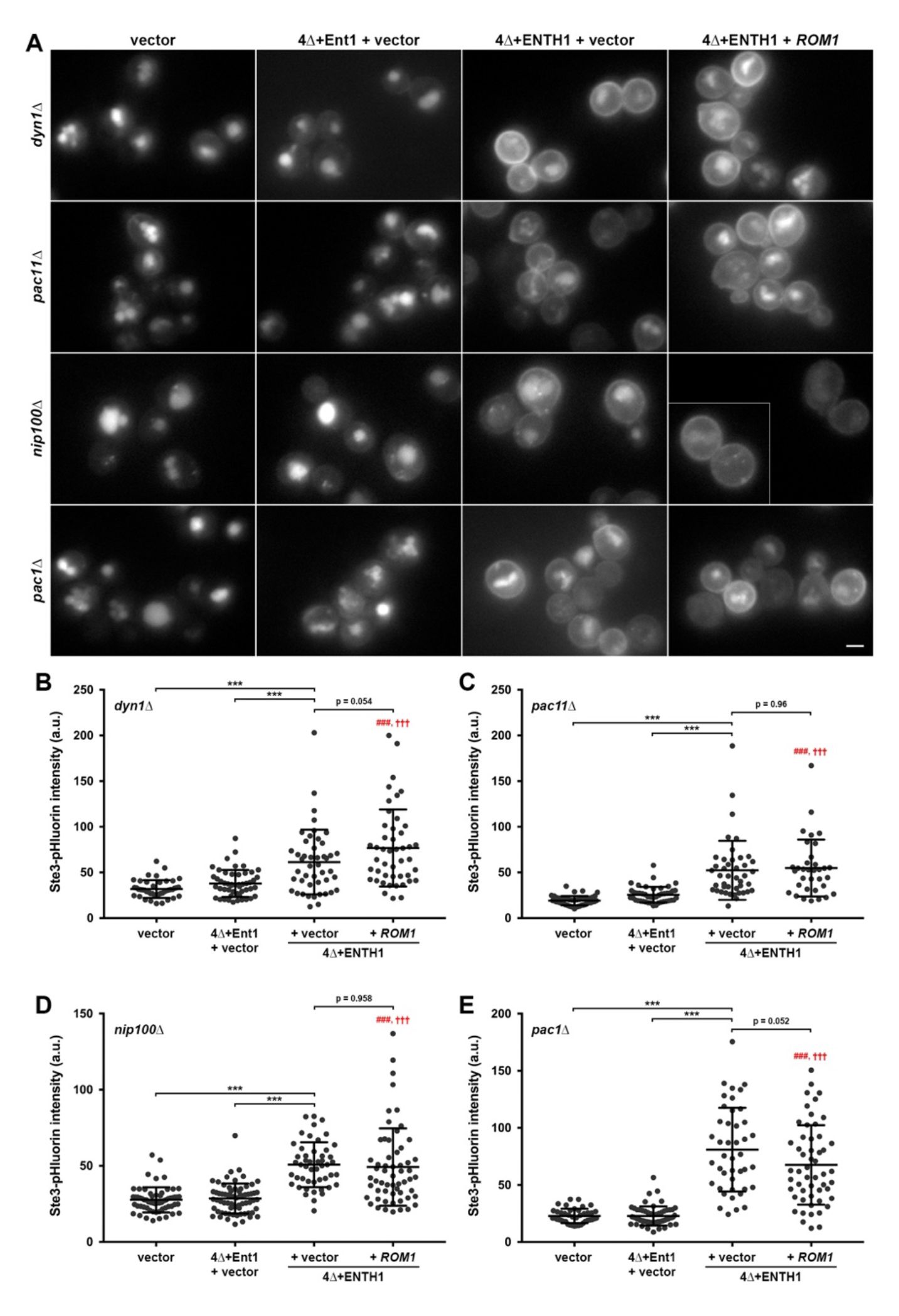
Requirement for dynein, dynactin, and microtubule plus-end tracking proteins in clathrin-independent endocytosis. Dynein (*dyn1*Δ and *pac11*Δ), dynactin (*nip100*Δ) and plus-end tracking (*pac1*Δ) mutants generated in WT (vector), 4Δ+Ent1 and 4Δ+ENTH1 backgrounds were transformed with vector or high-copy *ROM1* as indicated. (A) Cells expressing Ste3-GFP from the endogenous locus were imaged by fluorescence microscopy. Inset shows an additional cell at the same magnification. (B-E) Quantification of whole cell fluorescence intensity in cells expressing Ste3-pHluorin transformed as in *A*. (C) *dyn1*Δ (n=44, n=50, n=46 and n=46, respectively). (D) *pac11*Δ (n=47, n=45, n=43 and n=34, respectively). (E) *nip100*Δ (n=61, n=67, n=49 and n=58, respectively). (F) *pac1*Δ (n=42, n=59, n=43 and n=53, respectively). Mean ± s.d.; \*\*\**P*<0.001; ###*P*<0.001 compared to WT; †††*P*<0.001 compared to 4Δ+Ent1). Scale bar: 2 µm.

### Cortical microtubule capture is required for CIE in yeast

Once Myo2 delivers microtubule plus end to the cell cortex, an offloading event occurs in which the dynein intermediate chain Pac11 (in complex with microtubules) associates with Num1, a protein that forms stable patches at the cell cortex (Lee et al., 2003; Lee et al., 2005). These patches are sites of cMT anchoring, where Num1 binding to Pac11 leads to dynein activation by relieving the inhibitory activity of Pac1. Activated dynein is a minus end-directed motor, which pulls on the cytoplasmic microtubule to position the nucleus at the neck of large-budded cells in preparation for mitosis (Kahana et al., 1998). Since our data indicate that cytoplasmic microtubules and dynein/dynactin are involved in CIE, we generated *num1*Δ strains to test whether cMT anchoring is also required. As shown in in Fig. 7A, *num1*Δ did not affect vacuolar Ste3-GFP delivery in WT and 4Δ+Ent1 backgrounds transformed with empty vector, indicating that Num1 is not required for cargo internalization when CME is functional. In contrast, Ste3-GFP was prominently retained at the plasma membrane in *num1*Δ 4Δ+ENTH1 cells with empty vector, and this defective localization was not corrected by transformation with high-copy *ROM1*. Similarly, quantification of Ste3-pHluorin intensity in *num1*Δ cells showed low fluorescence in WT and 4Δ+Ent1 backgrounds with empty vector, in agreement with vacuolar localization seen for Ste3-GFP (Fig. 7B). As expected, Ste3-pHluorin intensity was significantly higher in *num1*Δ 4Δ+ENTH1 cells with empty vector. Transformation of *num1*Δ 4Δ+ENTH1 cells with high-copy *ROM1* failed to improve the elevated fluorescence intensity of Ste3-pHluorin compared to empty vector; in fact, *ROM1*-transformed cells were significantly brighter than the vector control in *num1*Δ 4Δ+ENTH1 cells, similar to *myo2^K1408A^* 4Δ+ENTH1 cells defective in Myo2-dependent microtubule transport (Fig. 2E). Thus, cMT capture and anchoring, in addition to transport of cytoplasmic microtubules, appears to be required for CIE.

**Figure 7:**
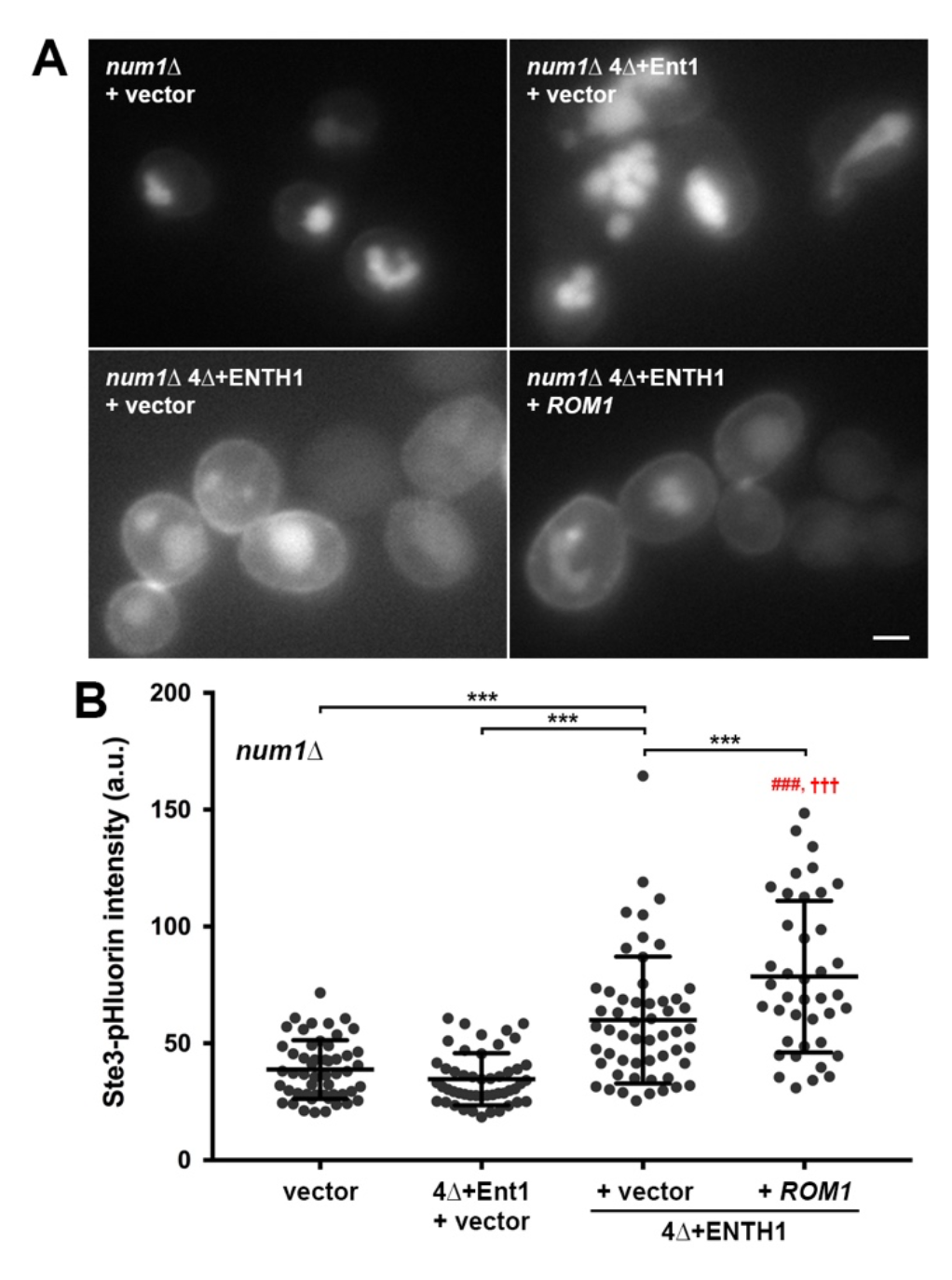
Requirement for the cortical microtubule capture protein Num1 in clathrin-independent endocytosis. *num1*Δ in WT (vector), 4Δ+Ent1 and 4Δ+ENTH1 backgrounds were transformed with vector or high-copy *ROM1* as indicated. (A) Cells expressing Ste3-GFP from the endogenous locus were imaged by fluorescence microscopy. (B) Quantification of whole cell fluorescence intensity in cells expressing Ste3-pHluorin transformed as in *A* (n=52, n=49, n=54 and n=40, respectively). Mean ± s.d.; \*\*\**P*<0.001; ###*P*<0.001 compared to WT; †††*P*<0.001 compared to 4Δ+Ent1). Scale bar: 2 µm.

In our examination of the role of microtubules and cMT capture machinery in CIE, we considered the possibility that mitotic spindle positioning may be altered in cells with defective CME or that strains with defective CIE may share a spindle orientation defect. Thus, we generated strains with a chromosomal integration of *GFP-TUB1*, expressed from the *TUB1* promoter, at the *LYS2* gene locus. WT, 4Δ+Ent1 and 4Δ+ENTH1 cells with empty vector or high-copy *ROM1* all had similar microtubule morphologies in unbudded and budded cells, with no obvious spindle misorientation, suggesting that inhibition of CME or overexpression of *ROM1* did not alter the microtubule cytoskeleton (Supplementary Fig. S2A). Additionally, we examined GFP-Tub1 in a variety of other strains harboring deletions in cMT transport and capture. With these strains, we observed spindle orientation similar to WT cells in some cases (*kar9*Δ, *bim1*Δ, *bik1*Δ and *pac1*Δ), but a severe spindle misalignment in cells with defective dynein, dynactin or cMT anchoring (*dyn1*Δ, *pac11*Δ, *nip100*Δ and *num1*Δ; Supplementary Fig. S2B). All spindle orientation phenotypes were similar when these mutations were generated in the 4Δ+Ent1 and 4Δ+ENTH1 backgrounds (data not shown). Our observations agree with numerous prior studies showing spindle misalignment in many mutants with defective cMT capture and nuclear positioning (Eshel et al., 1993; Farkasovsky and Küntzel, 1995; Kahana et al., 1998; Miller and Rose, 1998; Stuchell-Brereton et al., 2011). Notably, spindle misalignment is not necessarily required for a role of these proteins in CIE, since *kar9*Δ and *pac1*Δ, which had normal spindle orientation, both prevented Rom1-dependent activation of CIE in 4Δ+ENTH1 cells (Anderson et al., 2022; Lee et al., 2003). Moreover, we previously showed that Bni1 is required downstream of Rho1 for CIE, and we did not observe spindle orientation defects in *bni1*Δ 4Δ+ENTH1 cells (data not shown).

## Discussion

Although a variety of clathrin-independent endocytic pathways have now been observed in most eukaryotes, the molecular mechanisms governing CIE have remained elusive compared to our understanding of CME. Many aspects of CIE have not yet been addressed in detail, due in part to the relatively small number of proteins implicated in CIE and a lack of tools to study these pathways. In turn, this leaves open questions about (1) the degree of overlap in protein machinery between different CIE pathways or between CME and CIE, (2) the relative level of complexity required to generate clathrin-coated and clathrin-independent vesicles, (3) mechanisms for generating the force required for membrane deformation, curvature stabilization and vesicle scission in the absence of a clathrin coat, (4) how cells select and sort cargo into different endocytic pathways, and (5) the relative contribution of CME and CIE to cargo and membrane internalization.

Our previous discovery of the first CIE pathway observed in budding yeast has provided us with a genetically tractable system to begin answering these questions (Prosser et al., 2011). Earlier studies identified a signaling cascade involving the cell wall stress sensor Mid2, the Rho1 GEFs Rom1 and Rom2, the Rho1 GTPase and its effector Bni1 (along with Bni1-recruiting and - activating proteins in the polarisome) as central components of CIE in yeast. Moreover, the early-acting CME machinery protein Syp1 contributes to both clathrin-dependent and clathrin-independent internalization of cargos such as the di-and tri-peptide transporter Ptr2 (Apel et al., 2017). Syp1 contains an N-terminal Fes/CIP4 homology-Bin/Amphiphysin/Rvs (F-BAR) domain involved in sensing and inducing membrane curvature and a cargo-binding µ-homology domain (µHD) at its C-terminus, both of which are required for localization to and function at CME sites (Reider et al., 2009). Syp1 directly binds to and promotes internalization of Mid2, likely through CME since we have not observed Mid2 internalization under conditions where CIE promotes uptake of Ste3 or other cargos (D. Prosser, unpublished results); thus, it is unclear whether Syp1 plays mechanistically similar roles in CME and CIE. Finally, α-arrestins, which act as cargo-selective adaptors for the ubiquitin ligase Rsp5 during CME, also participate in CIE through interactions with proteins such as Rom2 and Rho1 (Prosser et al., 2015). Rsp5 and its interaction with α-arrestins is not required for CIE, even though the α-arrestins retain their cargo-selective roles in directing proteins to CIE. While the majority of yeast CME proteins tested to date are not required for CIE, these prior findings demonstrate that limited sets of proteins participate in both pathways.

The current study expands our understanding of CIE in yeast by identifying additional components of the pathway: the type V myosin Myo2, microtubules and plus end-binding proteins (Kar9, and possibly Bim1 and Bik1), regulators of cytoplasmic microtubule stability (Kip2 and Kip3), cMT capture (Num1), and dynein/dynactin (Dyn1, Pac11 and Nip100; Fig. 8). None of these proteins have been previously implicated in CME to our knowledge, and membrane trafficking roles for microtubules or microtubule-based motors remain largely undefined in yeast. Notably, tubulin and the plus end-tracking protein Bik1 were reported to play roles in trafficking of the v-SNARE Snc1, where mutations in Tub1 or Bik1 caused PM accumulation of Snc1 reminiscent of the endocytosis-defective Snc1^end-^ mutant (Boscheron et al., 2016; Lewis et al., 2000). Importantly, Snc1 mislocalization in *tub1-Glu* and *bik1*Δ mutants was corrected by expression of constitutively active Rho1^G19V^. This suggests a possible role for CIE in the observed effects, although contributions of CME cannot be ruled out. In our study, Bik1 was not required for CIE of Ste3-GFP (Fig. 3), which might be explained by overlapping functions with Bim1 or other components of microtubule plus end-binding or tracking. Alternatively, specific endocytic cargos may physically associate with different proteins involved in CIE and/or microtubule regulation to allow recruitment into an endocytic carrier, either directly or indirectly through proteins such as α-arrestins. For example, the α-arrestin Bul2 associates with the dynactin subunit Nip100, which may allow direct links between cargos and microtubule-based transport machinery (Wang et al., 2012). As we do not yet understand how cargo is selected into CIE pathways, identifying proteins that serve as CIE cargo-selective adaptors will be an interesting future direction.

**Figure 8:**
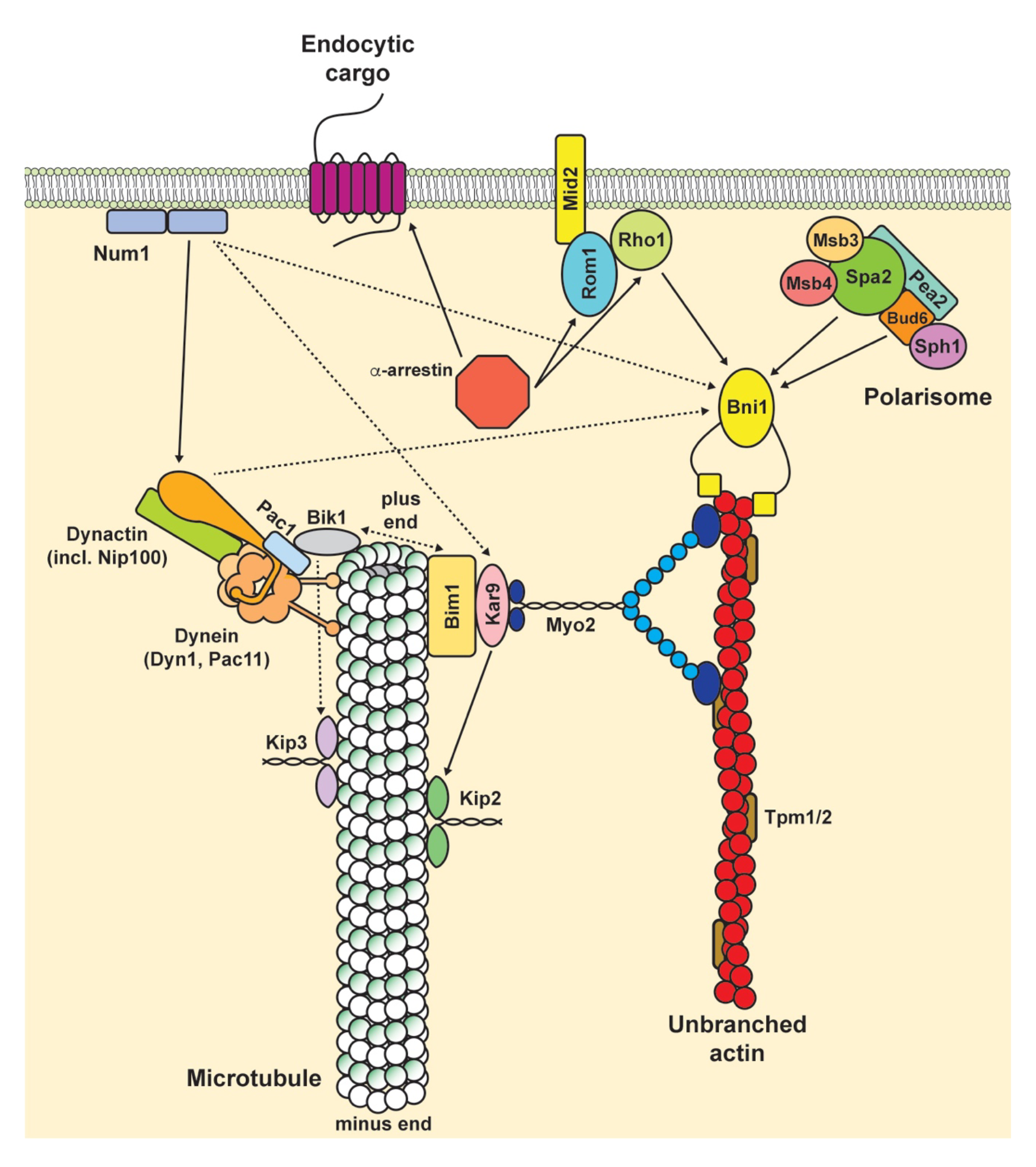
Model of protein modules involved in yeast clathrin-independent endocytosis. We previously identified several signaling modules and/or protein transport complexes that play roles in clathrin-independent endocytosis, including a cell wall stress-sensing module (Mid2, Rom1, Rho1), the polarisome (including Bni1, Bud6 and Spa2), unbranched actin filaments stabilized by tropomyosins (Tpm1 and Tpm2; Prosser et al., 2011), and α-arrestins (Prosser et al., 2015). Here, we add Myo2, proteins involved in microtubule plus-end transport (Kar9 and possibly Bim1), cytoplasmic microtubules and kinesin-related proteins involved in their stability (Kip2 and Kip3), dynein (including Dyn1 and Pac11), dynactin (including Nip100), microtubule plus-end tracking proteins (Pac1), and proteins involved in cortical microtubule capture (Num1) as additional factors that contribute to CIE. To date, none of these additional proteins have defined roles in clathrin-mediated endocytosis.

Separate from the role of Bik1 and tubulin in Snc1 trafficking, another study identified a role for the dynein light chain-family protein Tda2 in CME (Farrell et al., 2017). Rather than acting with microtubules and dynein, this novel role for Tda2 appears to rely on formation of a complex with Aim21 and the actin-capping proteins Cap1 and Cap2, leading to association with the CME protein Bbc1 and subsequent regulation of Arp2/3-dependent actin polymerization at cortical actin patches. Although these findings do not rule out a relationship between Tda2 and microtubules distinct from its contribution to CME, it is unlikely that Tda2 participates in CIE, since high-copy *ROM1* promotes Ste3-GFP internalization in *aim21*Δ 4Δ+ENTH1 cells (D. Prosser, unpublished results).

The degree of similarity between yeast and mammalian CIE remains a major unresolved question, in large part because of the limited set of proteins linked to any CIE pathway. As we add to the number of proteins involved in yeast CIE, it is tempting to speculate on how these relate to pathways in higher eukaryotes. Studies of several different pathways have identified roles for homologs of proteins involved in yeast CIE. For example, clathrin-independent ultrafast endocytosis occurs at synapses in mammals and *C. elegans*, and requires actin polymerization and formins (Bni1 in yeast; (Soykan et al., 2017; Watanabe et al., 2013a; Watanabe et al., 2013b)). Additionally, fast endophilin-mediated endocytosis (FEME) utilizes microtubules and dynein for inward transport of tubular endocytic carriers, as does clathrin-independent internalization of cholera toxin (Boucrot et al., 2015; Casamento and Boucrot, 2020; Day et al., 2015; Ferreira et al., 2021; Watanabe and Boucrot, 2017). Microtubules and the plus end-binding protein EB1 (Bim1 in yeast) contribute to endocytosis in *Drosophila* oocytes, although roles for clathrin in this process have not been addressed (Sanghavi et al., 2012). Lastly, RhoA and integrins (similar to yeast Rho1 and Mid2, respectively) are required for clathrin-independent compensatory endocytosis in bladder umbrella cells and for internalization of the IL-2 receptor (Khandelwal et al., 2010; Lamaze et al., 2001). While additional parallels likely exist, these suggest that the protein machinery involved in a variety of CIE pathways may at least partially overlap. Alternatively, yeast CIE may represent an ancestral form of clathrin-independent endocytosis that has branched or diverged during evolution. It will be interesting to test whether additional components of the yeast pathway contribute to CIE mechanisms in other eukaryotes, and our ability to identify additional CIE components using yeast genetics has the potential to rapidly expand our understanding of these pathways.

With an increase in our understanding of CIE pathways and the protein machinery that enables clathrin-independent internalization, insights into relationships between CIE and human diseases may also emerge. To date, homologs of several yeast CIE proteins have been linked to disease. For example, mutations in diaphanous-related formins, which are homologous to yeast Bni1, are associated with microcephaly and related developmental disorders (Labat-de-Hoz and Alonso, 2021). From our current study, Pac1 is related to the lissencephaly-related protein LIS1, while Kar9 is related to the APC protein involved in colorectal cancer (Geiser et al., 1997; Korinek et al., 2000). Moreover, the dynactin p150^Glued^ subunit associates with the huntingtin-associated protein HAP1 (Li et al., 1998); these may act in a complex involved in vesicular transport, particularly in axons. While direct roles for these proteins in mammalian CIE remain unclear, both in health and disease, it will be interesting to assess whether disease-linked mutations or losses of function alter cargo internalization, and whether such an effect might contribute to cellular dysfunction in disease.

Overall, our findings lead us to propose a model in which the interplay between myosin-and actin-dependent transport of microtubules, their cortical capture, and activation of dynein/dynactin promote CIE in yeast (Fig. 8). Myo2-dependent delivery of microtubule plus ends to the cell cortex requires on Kar9 and Bim1. As the plus end is transported through the cytoplasm, tracking proteins such as Bik1 and Pac1 promote loading and retention of dynein/dynactin, albeit in an inhibited state, and redundant pathways involving these proteins help to ensure the fidelity of microtubule delivery to the cell cortex (Adames and Cooper, 2000; Farkasovsky and Küntzel, 2001; Lee et al., 2000; Miller and Rose, 1998; Miller et al., 2000; Yin et al., 2000). Once delivered to the cell periphery, a hand-off event occurs wherein dynein associates with the cMT-capturing protein Num1, leading to cMT offloading from Myo2, dynein activation and minus end-directed motility (Lee et al., 2003; Lee et al., 2005). These events are thought to be necessary for the force generation required to position the nucleus at the bud neck during mitosis, thereby ensuring chromosome segregation and nuclear partitioning between mother and daughter cells. However, since dynein is tethered to Num1 at the cell cortex during this process, it seems plausible that minus end-directed motor activity could also exert inward force at the plasma membrane. Combined with Bni1-generated actin filaments and Myo2 motor activity, this could facilitate membrane bending in the absence of clathrin, which is needed to overcome the high turgor pressure in yeast cells (Aghamohammadzadeh and Ayscough, 2009). Our observation that cytoplasmic microtubules contribute to the Rho1 pathway supports a role in CIE, even though cytoplasmic microtubules appear to be sparse under most conditions in yeast. It is thus plausible that microtubule cytoskeleton-stabilizing environmental conditions could provide clues about how yeast CIE is regulated. Our future studies will examine this possibility, and will further test roles for the interaction network between actin, microtubules, and their respective motor proteins in membrane and protein internalization.

## Materials and Methods

### Yeast strains, plasmids, and growth conditions

Strains and plasmids used in this study are described in supplementary tables S1 and S2, respectively. Yeast were grown in liquid or plate-based YPD medium or on YNB (SD) medium lacking uracil, tryptophan, histidine and/or lysine for maintenance of non-essential plasmids. All cells were grown at 30°C and imaged at room temperature. PCR-based tagging of genomic loci and gene knockouts were performed as described previously (Goldstein and McCusker, 1999; Longtine et al., 1998). Transformations for plasmid uptake or for genomic integrations were performed using the LiAc method (Schiestl and Gietz, 1989). Unless otherwise specified, chemicals and reagents were purchased from Sigma-Aldrich or from Fisher Scientific.

### Construction and confirmation of MYO2 IQ repeat truncations

Plasmids for truncating Myo2 IQ repeats at the genomic *MYO2* locus with *HIS3* selection were generously provided by Dr. Anthony Bretscher (Cornell; (Schott et al., 1999; Schott et al., 2002)). The full-length control *MYO2^6IQ^* plasmid was linearized with SpeI, while the truncated *myo2^4IQ^* and *myo2^2IQ^* plasmids were linearized with BamHI, transformed into SEY6210 *MATα* wild-type or *ent1::LEU2 yap1802::LEU2* cells, and selected on YNB medium lacking histidine. *ent1::LEU2 yap1802::LEU2* cells with Myo2 modifications were then mated with *MATa ent2::HIS3 yap1801::HIS3* cells. Resulting diploids were transformed with pENT2.416 [*CEN URA3*], sporulated and tetrad dissected, and resulting *ent1::LEU2 ent2::HIS3 yap1801::HIS3 yap1802::LEU2*+pENT2.416 (4Δ+Ent2) cells were confirmed by PCR. Cells were then transformed with pENT1.414 or pENTH1.414 and grown on 5-FOA plates to select for loss of the pENT2.416 plasmid, yielding 4Δ+Ent1 and 4Δ+ENTH1 cells with wild-type or truncated Myo2. All Myo2 truncations were confirmed by PCR using the following primers: 5’- TTGATGGTGTTGTCTCAACTCAGAG-3’ and 5’-CATTGATTTGTGTAGCATTGACACC-3’, which amplify nucleotides 2047-3472 of the wild-type *MYO2* coding sequence containing the IQ repeat region. PCR products were then resolved by gel electrophoresis, and truncations were confirmed by differences in size compared to full-length *MYO2*.

### Construction and confirmation of myo2 cargo-binding domain mutant strains

Plasmids for low-copy expression of *MYO2* or *myo2^CBD^* mutants with *HIS3* selection were generous gifts from Dr. Lois Weisman (Univ. Michigan; (Catlett and Weisman, 1998; Eves et al., 2012; Ishikawa et al., 2003; Pashkova et al., 2006)). We used these as a starting point for generating chromosomally-integrated CBD mutants at the endogenous *MYO2* locus and expressed as the sole source of Myo2. To accomplish this, we isolated the 1.5 kb EcoRI fragment from wild-type and CBD mutant Myo2.413 plasmids containing the C-terminal CBD and 3’ untranslated region of *MYO2* for subcloning into the EcoRI site of pRS404. Resulting plasmids were confirmed by sequencing, and the orientation of all inserts placed the 5’ end of the Myo2 CBD proximal to the KpnI end of the polylinker. The plasmids were then linearized with NruI, transformed into SEY6210 *MATα* wild-type or 4Δ+pENT2.416 cells, and selected on YNB medium lacking tryptophan. 4Δ+Ent2 cells were then transformed with pENT1.317 or pENTH1.317 and selected on YNB medium lacking lysine. Cells were subsequently grown on 5- FOA plates to select for loss of the pENT2.416 plasmid, yielding Myo2 WT or CBD mutant-expressing 4Δ+Ent1 and 4Δ+ENTH1 strains. All WT and 4Δ strains with Myo2 CBD modifications were confirmed by isolation of genomic DNA and PCR amplification of the *MYO2* CBD using forward primer 5’-CTACCTCAAACACCATTAAAGGATG-3’ and T3 as the reverse primer. The resulting 1683 bp product spanning the 3’ end of *MYO2* upstream of the integration site through the 3’ untranslated region was then isolated using a PCR purification kit (Qiagen), and mutations were confirmed by sequencing using primer 5’- CTCATTTGTGGTGTTTGCTC-3’, which anneals to the *MYO2* CDS beginning at nucleotide 3777 downstream of the +1 site.

### Latrunculin A and Nocodazole treatment

Liquid cultures of WT, 4Δ+Ent1 and 4Δ+ENTH1 cells expressing Ste3-GFP and transformed with empty vector (pRS426) or high-copy *ROM1* were grown to mid-logarithmic phase at 30°C in YNB medium lacking uracil (Christianson et al., 1992; Ozaki et al., 1996). 0.7 OD_600_ of each strain was then pelleted by centrifugation at 3500 x g for 5 min, and cells were resuspended in 25 µl of YNB -ura medium supplemented with 200 µM Latrunculin A (Enzo Life Sciences), 15 µg/ml Nocodazole, or an equivalent volume of DMSO as a vehicle control. Cells were then incubated at 30°C for 2 h prior to imaging by fluorescence microscopy.

### Visualization of actin morphology

To visualize actin patches and cables, yeast cells were stained using a protocol modified from (Amberg, 1998). Briefly, *myo3*Δ *myo5*Δ cells were grown to mid-logarithmic phase in YNB medium lacking uracil for plasmid selection. Cells were fixed at room temperature for 30 min by addition of formaldehyde to a final concentration of 4% (v/v), then pelleted by centrifugation at 3000 x g for 5 min. Pellets were washed twice with phosphate-buffered saline (PBS) and stained for 1 h with Alexa 568 phalloidin (Thermo Fisher) dissolved in methanol, and diluted to a final concentration of 1.65 µM in PBS. Cells were then washed three times with PBS prior to imaging by fluorescence microscopy.

### Fluorescence microscopy and image analysis

Images were collected using either an Axiovert 200 inverted fluorescence microscope (Zeiss) equipped with a 100X, 1.4 NA Plan-Apochromat oil immersion objective, Sensicam (Cooke), X-Cite 120 PC light source, and SlideBook 4.2 software (3i) or using a DMi8 inverted fluorescence microscope (Leica) equipped with a 100X, 1.47 NA Plan-Apochromat oil immersion objective, LED3 fluorescence illumination system, Flash 4.0 v3 sCMOS camera (Hamamatsu) and LAS X v3.7.6.25997 (Leica). Within each experiment, all strains were imaged with the same acquisition parameters and on the same day. All imaging was performed using cells grown to mid-logarithmic phase.

Following acquisition, images were processed using Fiji/ImageJ2 v2.9.0/1.53t. Within each experiment, identical post-imaging processing was performed on all images to set identical minimum and maximum intensity levels, allowing direct comparison of protein localization and fluorescence intensity.

For visualization of phalloidin-labeled cells, z-stacks were collected at 0.25 µm step intervals spanning the entire depth of cells. Stacks were then collapsed into a maximum intensity z-projection image using Fiji/ImageJ2.

### Quantification of Ste3-pHluorin intensity

Quantification of Ste3-pHluorin intensity was performed as described previously (Prosser et al., 2010; Prosser et al., 2016). Briefly, random fields of cells for each condition analyzed were visualized by DIC prior to imaging by fluorescence microscopy. All conditions within an experiment were imaged on the same day, using identical acquisition parameters. Background subtraction was then performed on 16-bit images, individual cells were selected for measurement of whole-cell fluorescence intensity, and values were corrected for cell size. Pre-determined criteria for exclusion of cells from analysis are described in (Prosser et al., 2016).

### Statistical analysis

Power analysis was performed to determine population sizes using G*Power v3.1.9.6, with type I error α=0.05, type II error β=0.2 (power, 1-β=0.8), and a moderate effect size f=0.3. For experiments with four groups (WT, 4Δ+Ent1 and 4Δ+ENTH1 backgrounds with empty vector; 4Δ+ENTH1 with high-copy *ROM1*) using one-way ANOVA, the analysis recommended a minimum of 32 cells measured per condition.

Statistical significance for all quantitative experiments was assessed using one-way ANOVA followed by Tukey’s Multiple Comparison test in Prism 7 (GraphPad).

## Acknowledgements

We would like to gratefully acknowledge gifts of *MYO2* plasmids from Drs. Lois Weisman (University of Michigan) and Anthony Bretscher (Cornell). We also thank Drs. M. Andrew Hoyt (Johns Hopkins), Trina Schroer (Johns Hopkins) and Lois Weisman, as well as members of the Wendland and Prosser labs for helpful discussions, Joanna Poprawski and Lydia Nyasae for excellent technical assistance, and J. Michael McCaffery and Erin Pryce (Johns Hopkins Integrated Imaging Center) for advice on imaging. This work was initiated in the laboratory of Dr. Beverly Wendland (Johns Hopkins), and her advice, support, and mentorship have been immeasurably valuable; we acknowledge her contributions with the strongest possible gratitude.

